# Cystathionine β-synthase gene inactivation dysregulates major urinary protein biogenesis and impairs sexual signaling in mice

**DOI:** 10.1101/2022.07.10.499408

**Authors:** Ewa Bretes, Jacek Wróblewski, Monika Wyszczelska-Rokiel, Hieronim Jakubowski

## Abstract

Reproductive success in mice depends on major urinary proteins (Mup) that facilitate sexual interactions between females and males. Deletion of cystathionine β-synthase (Cbs) gene, a metabolic gene important for homeostasis of one-carbon metabolism, impairs reproduction by causing female infertility in mice. Here we examined Mup biogenesis and sexual signaling in *Cbs*^−/−^ *vs*. *Cbs*^+/−^ mice. We found that total urinary Mup protein was significantly reduced in male and female *Cbs*^−/−^ *vs. Cbs*^+/−^ mice. SDS-PAGE/Western blot, ESIMS, and RT-qPCR analyses of the liver, plasma, and urinary proteins identified a male-specific Mup20 in *Cbs*^−/−^, but not *Cbs^+/−^* females. As other Mups were significantly reduced, the 18,893 Da Mup20 became the most abundand in urine of *Cbs*^−/−^ females and males. Effects of *Cbs* genotype on 18,645 Da, 18,693 Da, and 18,709 Da Mup species abundance were Mup and sex-specific. *Cbs*-dependent changes in hepatic Mups and Mup20 expression were similar at the protein and mRNA level. Changes in Mups, but not in Mup20, can be explained by downregulation of hepatic Zhx2 and Ghr receptors in *Cbs*^−/−^ mice. Behavioral testing showed that *Cbs*^+/−^ females were attracted to *Cbs*^+/−^ but not to *Cbs*^−/−^ male urine. *Cbs*^+/−^ males did not countermark urine of *Cbs*^−/−^ males but countermarked urine of other *Cbs*^+/−^ males and were attracted to urines of *Cbs*^−/−^ as well as *Cbs*^+/−^ females. *Cbs*^−/−^ males did not countermark urine of *Cbs*^+/−^ males but were still attracted to urines of *Cbs*^+/−^ females. Taken together, these findings show that *Cbs*, a metabolic gene, plays an important role in the regulation of Mup biogenesis and sexual signaling in mice.

---

In humans and other mammals, homocysteine (Hcy), a sulfur-containing non-protein amino acid, originates from the dietary protein methionine [1]. Subsequent metabolic conversion of Hcy to cysteine and methionine is important for maintaining the homeostasis of redox status and folate metabolism. Because of these interconnections, dysregulated Hcy metabolism, leading to hyperhomocysteinemia (HHcy), is associated with cardiovascular [2] and neurological diseases [3, 4], cancer [5–7] and reproductive impairment [8, 9]. In humans, severe HHcy due to cystathionine β-synthase (CBS) deficiency [10] results in 26-48% fetal loss [11]. Mild HHcy is also a risk factor for pregnancy complications such as early pregnancy loss [12], preeclampsia [13], neural tube defects [8,14] as well as for male infertility [15]. In mice, *Cbs*^−/−^ females are infertile [16] due to altered estrus cycle and uterine failure [17], impaired decidualization and altered gene expression in uterus implantation sites [18].

Sexual behavior and reproduction in mice is governed by nonvolatile and volatile pheromones excreted in the urine [19]. Nonvolatile pheromones, known as major urinary proteins (Mups), are the products of the most highly expressed genes in the liver (5% of total mRNA), which is their main source [20]. Mups are a family of 18-19 kDa proteins encoded by the gene cluster on mouse chromosome IV. The genome of the C57BL/6J mice contains about 20 *Mup* genes, with only some expressed and excreted in urine. The products of *Mup* genes have an extremely high sequence similarity of >97%. The expression of Mups is a sexually dimorphic trait, with male mice excreting in their urine more Mups than females. Males increase scent marking in response to females or female scent [21]. Mups determine the sexual attractiveness of male urinary odor to females and exposure to male urine affects female’s behavior. Increased Mups excretion is correlated with male, but not female, reproductive success. Mups mediate female’s maternal aggression [22] and male-male territorial aggression [23]. Mup20, expressed exclusively in males, is inherently attractive to females, and stimulates their associative learning by evoking persistent remembered attraction and physical location of the male scent marks [24, 25].

A variety of factors can affect mouse sexual behavior and the type and amount of Mup excreted in the urine [19]. These include age, social status and interactions, nutrition and health. However, how cellular metabolic processes, known to be important for the reproductive success in mice, affect Mup expression and sexual behavior is not known. For this reason, we have studied factors affecting Mup expression at the level of protein and mRNA in *Cbs*^−/−^ *vs*. *Cbs*^+/−^ female and male mice. We also examined sexual preferences in these mice by studying their responses to urine scent marks from *Cbs*^−/−^ male and female mice. One of the unexpected findings was that the expression of the exclusively male Mup20, absent in *Cbs*^+/−^ female mice, was de-repressed in *Cbs*^−/−^ females.

## MATERIALS AND METHODS

### Mice

Transgenic *Tg-I278T Cbs*^−/−^ mice on the C57BL/6J background [26] were housed and bred at the Rutgers-New Jersey Medical School Animal Facility. Transgene harboring human *CBS I278T(Tg-I278T)* variant is used to rescue the neonatal lethality associated with homozygosity for *Cbs*^−/−^. In these animals, the human mutant CBS, which has less than 3% of the activity of wild-type enzyme, is under control of the zinc-inducible metallothionein promoter, which allows to rescue the neonatal lethality phenotype of *Cbs*^−/−^ in mice by supplementing the drinking water of pregnant dams with 25 mM zinc chloride. Zinc-water is replaced by plain water after weaning at 30 days. Only *Cbs*^−/−^ individuals exhibit changed phenotype characterized by facial alopecia, thin, smooth, shiny tail, reduced body weight and shortened life span compared to *Cbs*^+/−^ [26, 27]. Mouse *Cbs* genotypes were established by PCR using the following primers: forward 5□-GGTCTGGAATTCACTATGTAGC-3□, wild type reverse CGGATGACCTGCATTCATCT-3□, mutant reverse: GAGGTCGACGGTATCGATA-3□. The mice were fed with a standars rodent chow (LabDiet5010; Purina Mills International, St. Louis MO, USA). Two-to 12-month-old mice were used in experimnets. Control animals were *Cbs*^+/−^ siblings. Animal procedures were approved by the Institutional Animal Care and Use Committee at Rutgers-New Jersey Medical School.

### Urine collection and Mup protein quantification

Urine samples were collected by holding mice by the scruff and tail base over the clean petri dish. If necessary, the bladder was gently massaged. Urine was immediately transferred to a eppendorf tube, frozen in liquid nitrogen, and stored at-70 °C. Total urinary protein concentration was determined using the tannin turbidimetric assay [54]. Briefly: 100 μl of tannin reagent (10 g tannin dissolved in 98 ml 1N HCl and 2 ml of phenol) was added to 100 μl appropriately diluted urine and incubated at 30°C on a mixer set to 150 rpm. After 10 min, 150 μl of 0.1 % arabic gum solution was added and turbidity was measured at 595 nm using BioTek microplate reader ELx808.

### Liver and plasma collection

Blood was collected from cheek veins into Eppendorf tubes containing 1% (v/v) 0.5 M EDTA. After centrifugation (2000× g, 10 min, 4°C), separated plasma and cells were frozen at −80°C. Mice were euthanized using CO_2_ and the organs collected and frozen on dry ice. Livers were pulverized with dry ice using a mortar and pestle and stored at −80°C. Proteins were extracted from powdered tissues with RIPA buffer containing protease inhibitors cocktail (MilliporeSigma), added at a ratio 10 μL/1 mg of the pulverized liver with sonication (Bandelin SONOPLUS HD 2070) on wet ice (five 30-s strokes with 1 min cooling interval between strokes). Extracts were clarifed by centrifugation (15,000 g, 5 min, 4°C).

### Western blots of Mup and Zhx2 proteins

Liver proteins were separated by SDS-PAGE on 14 % gels (2.5 μg protein/lane for MUP analysis) or 10% gels (20 μg protein/lane for Zhx2 analysis). Proteins were transferred to PVDF membrane (Bio-Rad) for 20 min at 0.1 A, 25 V using Trans Blot Turbo Transfer System (Bio-Rad). After blocking with 5 % non-fat dry milk in TBST buffer (1 h, room temperature), the membranes were and incubated with anti-Mup1 (abcam95198), anti-Zhx2 (abcam 205532) for 1 hour, and with anti-Gapdh (Sigma-Aldrich PLA0125) antibodies for another hour. Membranes were washed three times with TBST buffer, 10 min each, and incubated with goat anti-rabbit IgG secondary antibody conjugated with horseradish peroxidase. Positive signals were detected using Western Bright Quantum-Advansta K12042-D20 and GeneGnome XRQ NPC chemiluminescence detection system. Bands intensity was calculated using Gene Tools program from Syngene.

### ESI-MS

Urine samples from *Cbs*^−/−^ and control mice were analyzed in a proteomic facility at the Institute of Biochemistry and Biophysics, Polish Academy of Sciences, Warsaw, Poland. Samples of dried, filtered mouse urine were dissolved in 20 μL of 0.1% TFA in water; 2 μL ofthe resulting solution were added to 500 μL of ACN:H2O:HCOOH (50:50:0.3 %). One fifth of such solution (100 μL) was used per assay. Analyses were performed on QToF Premier mass spectrometer (Waters/Micromass, Manchester, UK). The source ES parameters were as follows: voltage on the ESI probe tip (capillary voltage) 2.5 kV, sampling cone 30 V, extraction cone 2.0 V, ion guide 3.8 V, source block temperature 80°C, desolvatation temperature 40°C, glas cone 0 L/hr, desolvatation 650 L/hr, syringe pump flow rate 10 μL/min. The time span of one scan was 1 s, and ~200 scans were accumulated. Positive ion spectra were decon-voluted using the MaxEnt module of the MassLynx suite (Micromass).

### RNA isolation, cDNA synthesis, RT-qPCR analysis

Total RNA was isolated from mouse liver using PureLink RNA mini KIT (Thermo Fisher Scientific). cDNA synthesis was carried out using High-Capacity cDNA Reverse Transcription Kit (Applied Biosystems) according to manufacturer recommendation. Nucleic acid concentration was measured using NanoDrop (Thermo Fisher Scientific). Quantitative PCR was performed with SYBR Green Mix and CFX96 thermocycler (Bio-Rad).: 2^(-ΔΔCt)^ method was used to calculate the relative expression levels [28]. Data analysis was performed with the CFX Manager™ Software, Microsoft Excel, and Statistica. RT-qPCR primer sequences are listed in **Table S1.**

### Behavioral testing

The testing procedure was adopted from ref. [29]. Wild type C57BL/6J female mice (n = 12), male *Cbs*^−/−^ mice (n = 7) and their male *Cbs*^+/−^ littermates (n = 16) on the C57BL/6J background, six-to seven-month-old, were tested individually in separate cages in the morning in a period from April to June. Each mouse was teste in a cage of the same dimensions as the home cage. The bottom of each cage was covered with a sheet of Schleicher & Schuell GB002 gel blot paper (25.5 x 15 cm, 0.4 mm thick). The sheet was divided into three 8.5-cm sections with two faint pencil marks. Tested urine was spotted in two 8-μL aliquots 4 cm apart in the center of one lateral region. Two 8-μL aliquots of water were spotted in the opposite lateral region as a control. The central region was left clean. Under dim light, a mouse was placed in the central region and left free to move for 30 min. At the end of experiment, the paper was removed from the cage and a grid of 1-cm squares was drawn with faint pencil marks in the two lateral regions of the paper. The number of urine drops, voidings, and area covered by urine were counted under UV light. Each mouse was tested 2-4 times, each test separated by up to 12 days, and an average response for each mouse was calculated.

### Statistical analysis

The results were calculated as mean ± standard deviation. A parametric unpaired t test was used for comparisons between two groups of variables, p<0.05 was considered significant. Statistical analysis was performed using Statistica, Version 13 (TIBCO Software Inc., Palo Alto, CA, USA, http://statistica.io).

## RESULTS

### Sex-independent effect of *Cbs* genotype on Mup excretion in urine

We found that Mup levels were significantly reduced (2-fold) in 4-6-month-old *Cbs*^−/−^ *vs. Cbs*^+/−^ male mice (7.7 vs. 14.2mg/mL, *P* = 0.009) (Figure 1A). Mup concentration was also similarly reduced (2-fold) in *Cbs*^−/−^ *vs. Cbs*^+/−^ female mice (1.4 vs. 3.1 mg/mL, *P* = 0.0001). Mup concentration was significantly lower in *Cbs*^+/−^ female compared to *Cbs*^+/−^ male mice. Similar reduction in Mup levels was als found in *Cbs*^−/−^ female compared to *Cbs*^−/−^ male mice. These findings indicate that the influence of *Cbs*^−/−^ genotype on Mup levels is sex-independent.

**Figure 1.**
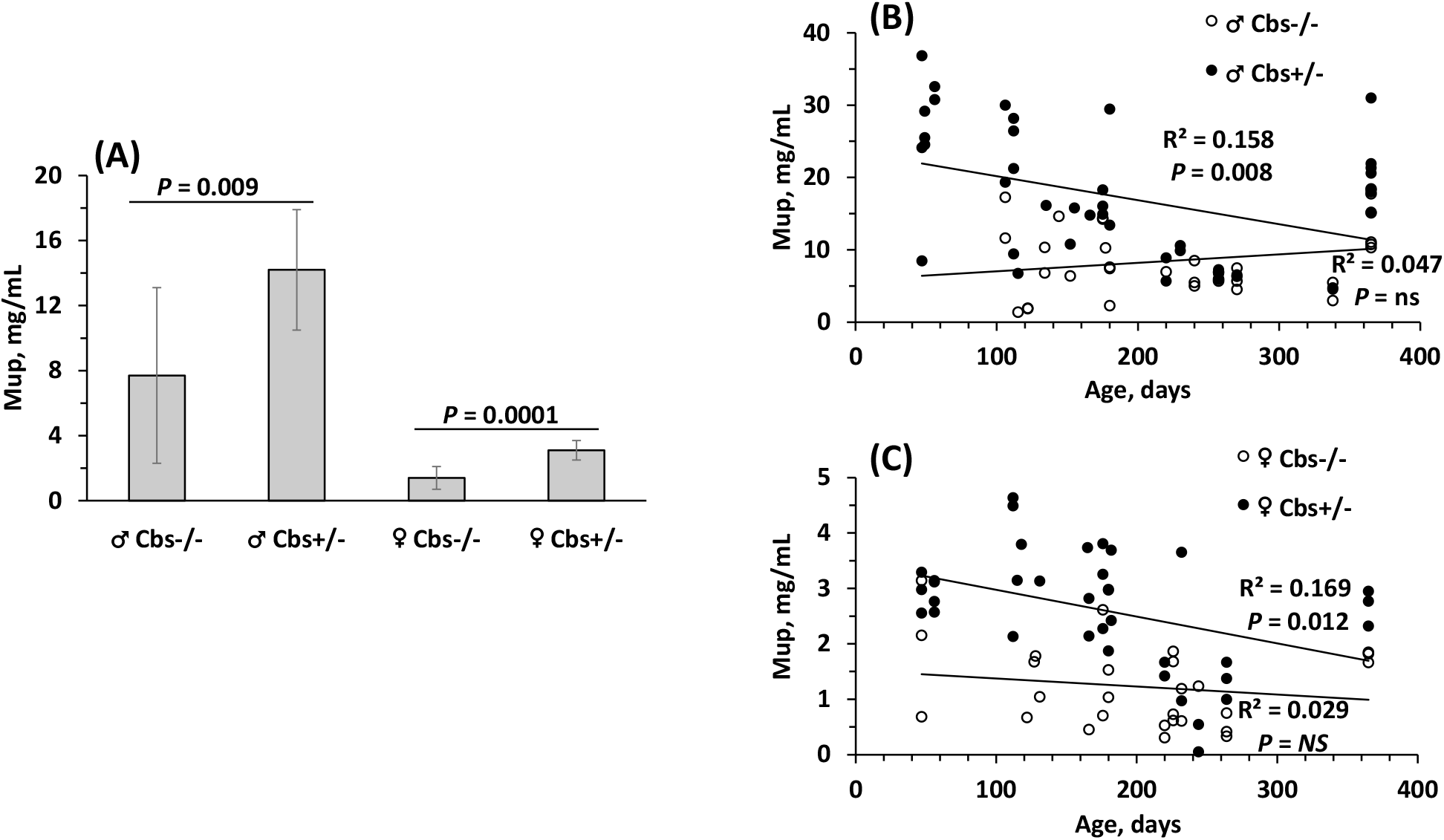
Total Mup levels were significantly reduced in *Cbs*^−/−^ *vs. Cbs*^+/-^ mice. (A) Mup levels were quantified by measurements of protein content in urines from 4- to 6-month-old □ *Cbs*^−/−^ (n = 11), □ *Cbs*^−/−^ (n = 9) and control □ *Cbs^+/-^* (n = 8) and □ *Cbs^+/-^* (n = 9) mice. (B, C) Effects of age on Mup levels were abrogated in *Cbs*^−/−^ mice. Total Mup protein was quantified in urines from mice aged from 47 to 365 days: (B) □ *Cbs*^−/−^ (n = 33) and control □ *Cbs*^+/−^ (n = 43); (C) □ *Cbs*^−/−^ (n = 27) and control □ *Cbs*^+/−^ (n = 36). *NS*, nor significant.

### *Cbs* genotype abrogates effects of age on Mup levels

To examine effects of age on Mup, we quantified Mup levels in urines from male *Cbs*^−/−^ (n = 33), male *Cbs^+/-^* (n = 43), female *Cbs*^−/−^ (n = 27), and female *Cbs^+/-^* (n = 36) mice aged 47 to 362 days.

In bivariate correlation analyses, we found that Mup levels were not affectd by age in *Cbs*^−/−^ male mice (*P* = 0.225, **Figure 1B**). Hovever, the dependence of Mup levels on age was observed in control *Cbs*^+/−^ males, as expected (*P* = 0.008). Similar loss of the age-Mup relationship was observed in *Cbs*^−/−^ females (*P* = 0.408 vs. *P* = 0.012 for *Cbs*^+/−^females, **Figure 1C**).

In multiple regression analyses in a model including Mup levels as a dependent variable and *Cbs* genotype, sex, and age as in dependent variables, *Cbs* genotype (β = −0.29, *P* = 0.000; lower in *Cbs*^−/−^ mice), sex (β = 0.68, *P* = 0.000; higher in males), and age (β = −0.15, *P* = 0.012; lower in older mice) were significant determinnats of Mup; adjusted R^2^ was 0.54, *P* = 0.000 **(Table 1)**.

**Table 1.**
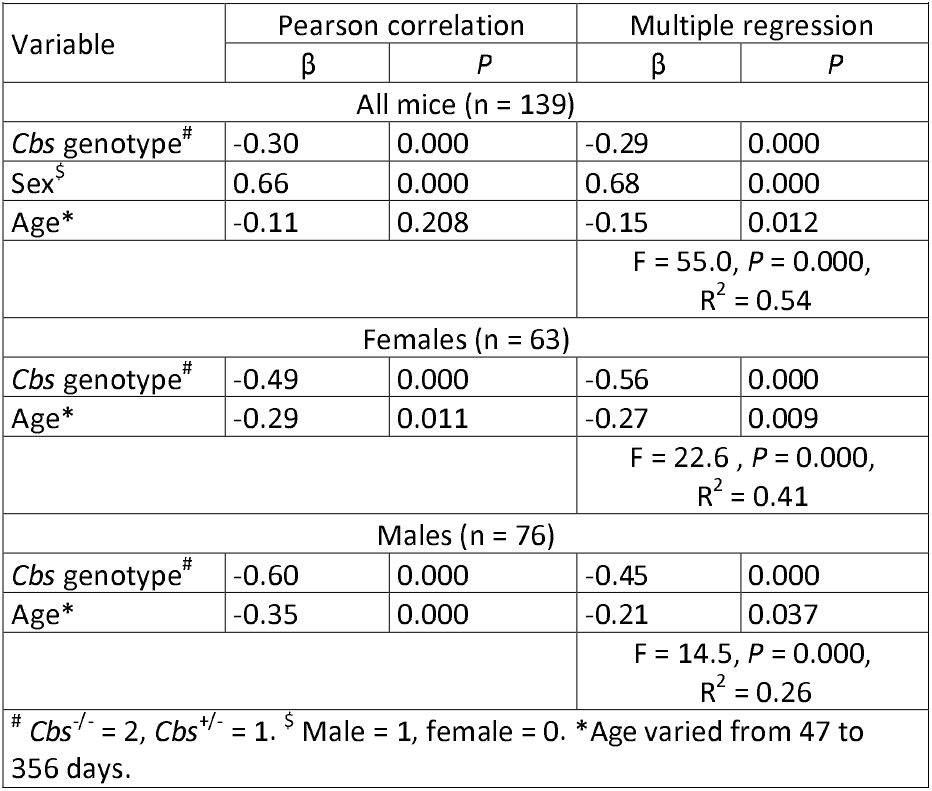
Determinants of urinary Mup levels in mice.

Multiple regression analysis stratified by sex showed that *Cbs* genotype and age were strong determinants of Mup levels both in female and male subgroups (**Table 1**).

### Mup20 is excreted in urine of *Cbs*^−/−^ females

To identify *Cbs-* genotype dependent changes in the levels of individual Mup proteins, we carried out SDS-PAGE and ESI-MS analyses of mouse urine. SDS-PAGE analyses showed that the intensity of Mups bands were reduced in *Cbs*^−/−^ *vs. Cbs*^+/-^ mice, both for females and males **(Figure 2A).** Remarkably, the male Mup20 band, absent in control female *Cbs*^+/-^ mice, was present in urines of female *Cbs*^−/−^ mice **(Figure 2).** In contrast in males, the presence of Mup20 was not affected by *Cbs* genotype.

**Figure 2.**
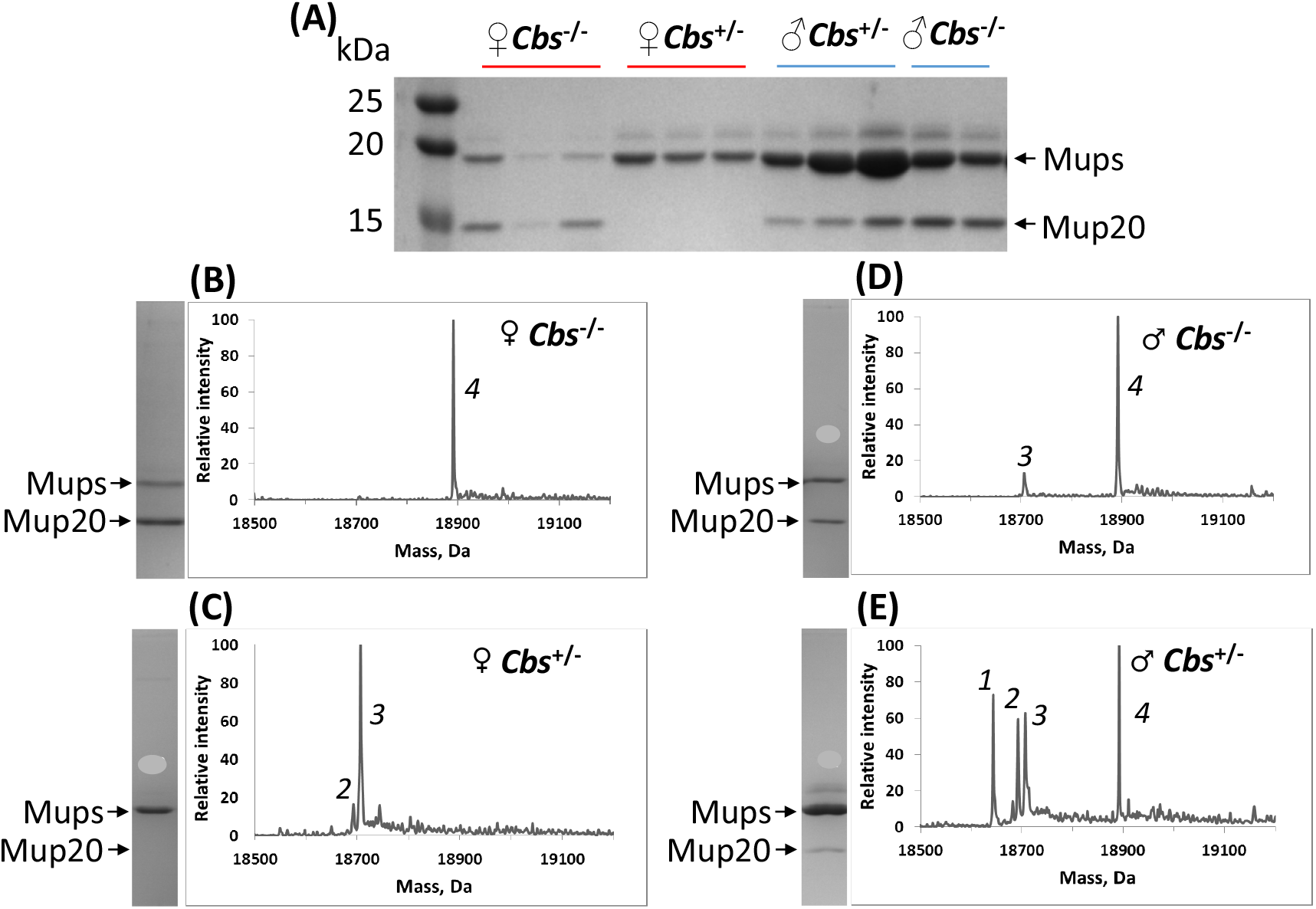
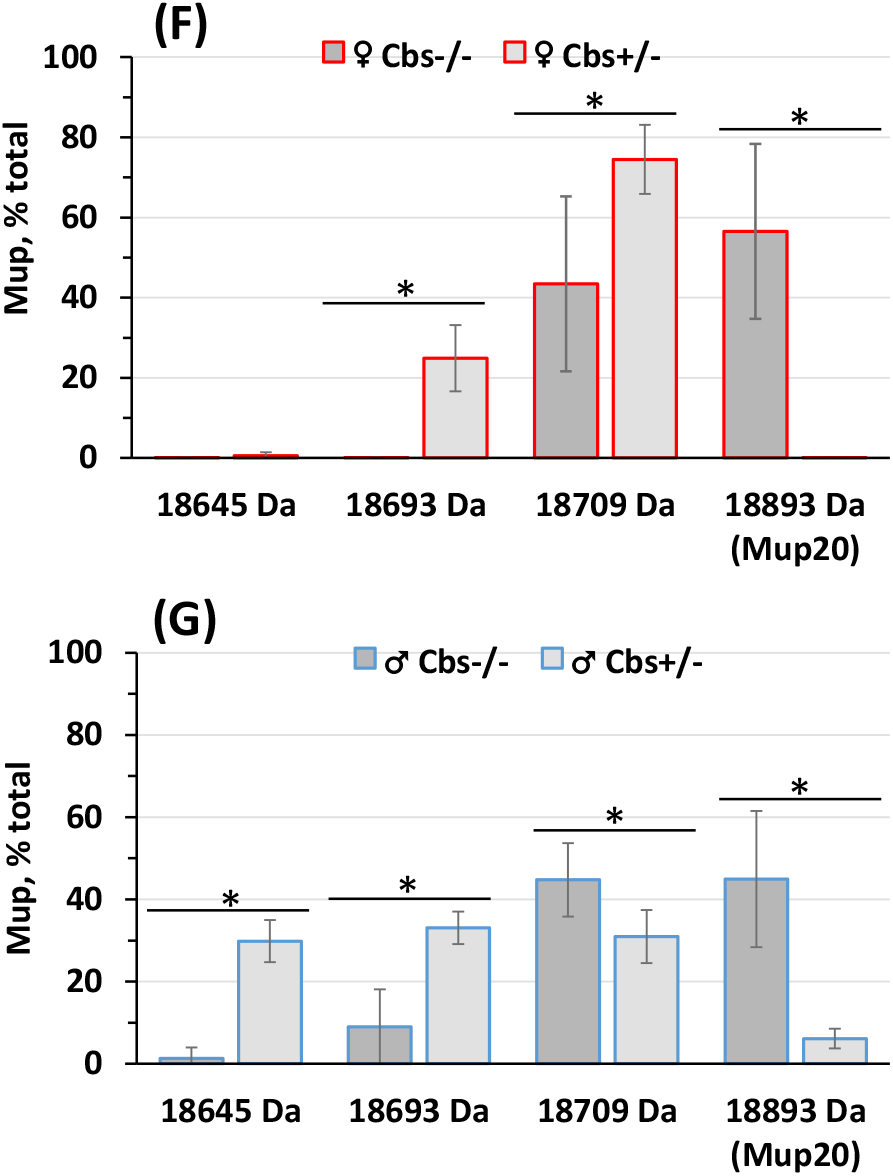
Male-specific Mup20 is present in urine of female *Cbs*^−/−^ mice. (A) Mup proteins from mouse urine were separated into to individual components by SDS-PAGE and visualized by stating with Coomassie Blue. (B, C, D, E) ESI-MS profiles of individual Mup proteins in urines from □ *Cbs*^−/−^ (B), □ *Cbs^+/-^* (C), □ *Cbs*^−/−^ (D), and □ *Cbs*^+/−^ (E) mice. Pictures from corresponding SDS-PAGE analyses are shown to the left of each ESI-MS panel. Molecular weights of numbered peaks are: 1,18,645 Da; 2, 18,693Da; 3,18,709 Da; 4,18,893 Da. The 18,893 Da protein is Mup20. (F, G) Abundance of individual Mup species in *Cbs*^−/−^ and *Cbs*^+/−^ mice. (F) □ *Cbs*^−/−^ and □ *Cbs*^+/−^ mice; 5 animals/group. (G) □ *Cbs*^−/−^ and □ *Cbs*^+/−^ mice; 5 animals/group. A peak intensity for each Mup specie was divided by the intensity for all peaks, expressed as % of total for each mouse, and averaged for each group of five animals. An asterisk ‘*’ indicates a significant difference.

ESI-MS analyses showed that just one Mup species of 18,893 Da (peak 4), correponding to Mup20, was present in urine from *Cbs*^−/−^ female **(Figure 2B).** Mup20 was absent in control *Cbs^+/-^* female **(Figure 2C)** but was present in *Cbs*^−/−^ and *Cbs^+/-^* male **(Figure 2D, E).** Urine from the control *Cbs^+/-^* male showed three more Mup species of 18,643.5 Da, 18,692.5 Da, and 18,706.5 Da. Control *Cbs^+/-^* female urine showed two middle Mup proteins of 18,692.5 Da and 18,706.5 Da. Similar results were obtained with five additional mice per each sex and *Cbs* genotype group **(Figure S1).**

### Sex-specific effects of *Cbs* genotype on the urinary abundance of individual Mup species

Quantification of the four major Mup species in *Cbs*^−/−^ and *Cbs*^+/−^ mice from the ESI-MS experiments is shown in **Figure 2F** for female mice and in **Figure 2G** for male mice. In control *Cbs^+/-^* female urines, two Mup species of molecular weight 18,709 Da and 18,693 Da, accounting for 75.5% and 24.5% of total were present. Strikingly, in *Cbs*^−/−^ female urines, a new peak corresponding to 18,893 Da (Mup20) appeared and the 18,693 Da Mup disappeared. The 18,893 Da (Mup20) and 18,709 Da Mup species accounted for 56.5% and 43.5% of total. The 18,709 Da Mup, present in *Cbs*^+/−^ female urines, was significantly downregulated in *Cbs*^−/−^ female urines **(Figure 2F).**

In control *Cbs*^+/−^ male urines, four Mup species of molecular weight 18,645 Da, 18,693 Da, 18,709 Da, and 18,893 Da (Mup20), accounting for 29.9%, 33.1%, 30.9%, and 6.1% of total were present. In *Cbs*^−/−^ male urines, 18,709 Da and 18,893 Da (Mup20) species were significantly upregulated and became dominant, representing 44.8% and 45.0% of total. In contrast, the 18,645 Da and 18,693 Da Mup species were significantly downregulated and represented 1.3% and 9.0% of total in urines of *Cbs*^−/−^ males **(Figure 2G).**

### Effects of *Cbs*^−/−^ genotype on liver and plasma Mup levels

To assess which stage of Mup biogenesis and transit to urine is affected by the *Cbs* genotype, we examined Mup expression and abundance in the liver and plasma. We quantified liver and plasma Mup proteins by SDS-PAGE and Western blotting using anti-Mup antibodies that recognizes Mup20 and othre Mups. Western blot analyses showed that Mup20 was present in the liver **(Figure 3A)** and plasma **(Figure 3B)** of *Cbs*^−/−^ females, *Cbs*^−/−^ males, and *Cbs*^+/−^ males, but was absent in liver **(Figure 3A)** and plasma **(Figure 3B)** of *Cbs*^+/−^ females.

**Figure 3.**
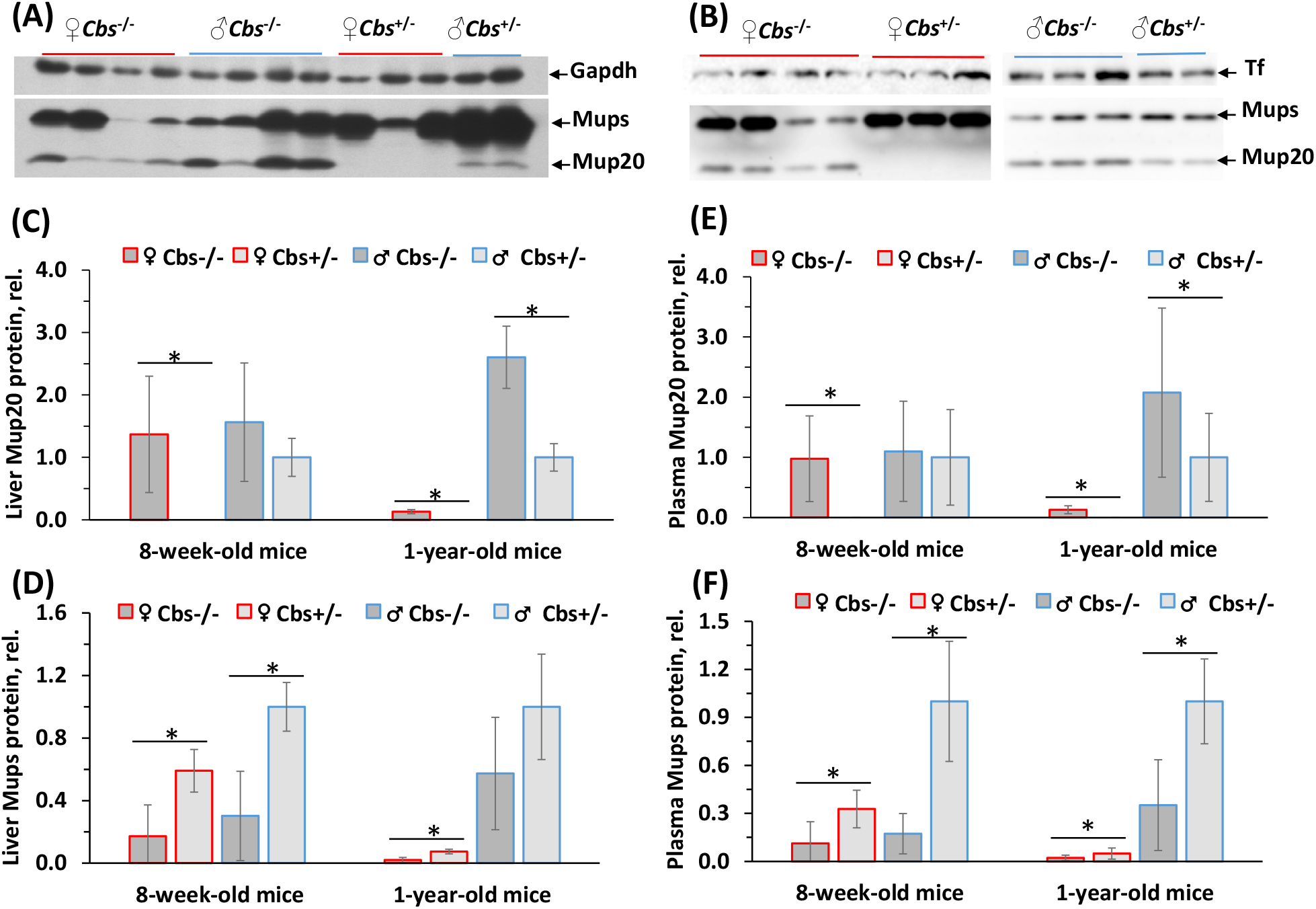
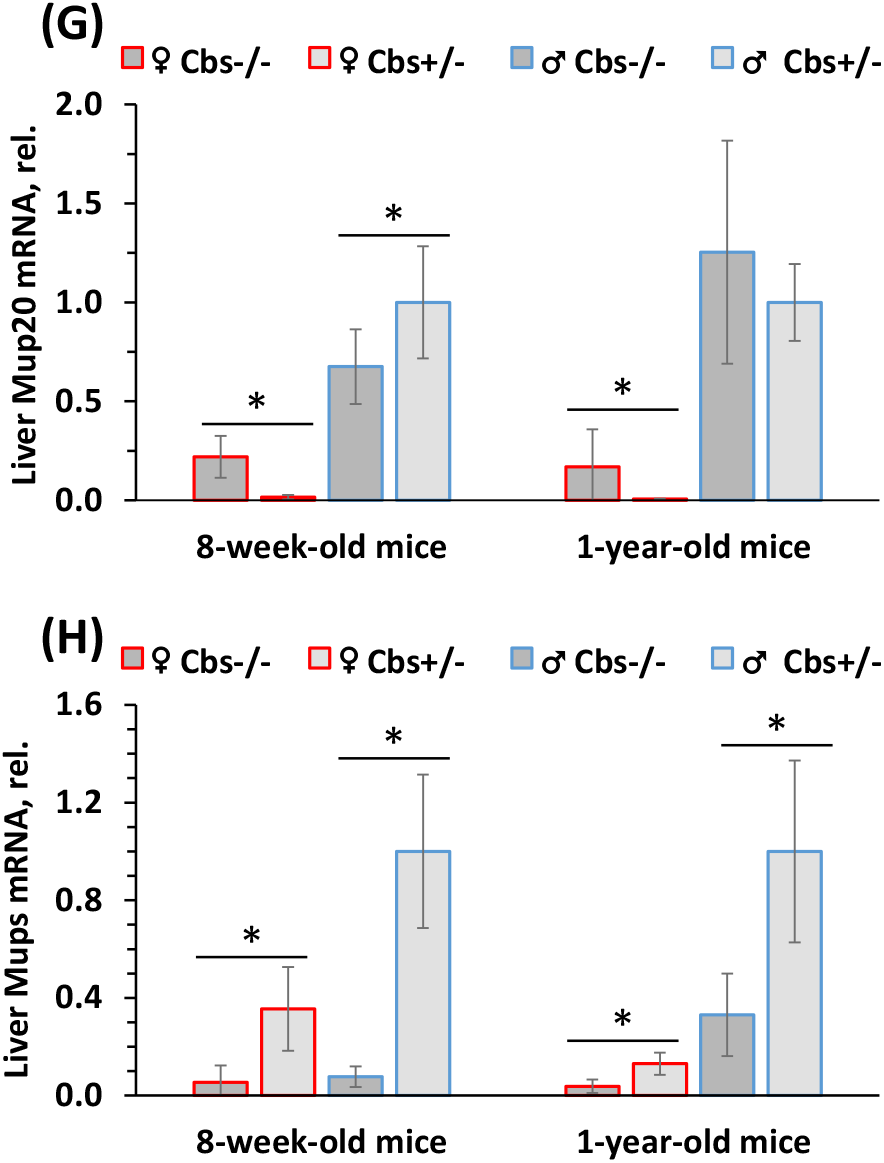
Male-specific Mup20 is expressed in the liver and present in plasma of □ *Cbs*^−/−^ mice. Liver and plasma proteins were separated by SDS-PAGE and Mup proteins were visualized by Western blotting using anti-Mup antibody. Gapdh was used as a reference for liver Mups; transferin (Tf) was used as a reference for plasma Mups. (A, B) Representative Western blots of liver (A) and plasma (B) proteins. (C, D, E, F) Quantification of Western blots of liver Mup20 (C) and Mups (D), plasma Mup20 (E) and Mups (F) in □ *Cbs*^−/−^, □ *Cbs*^+/−^, □ *Cbs*^−/−^, and □ *Cbs*^+/−^ mice; plasma 3-7 mice/group, liver 5 mice/group. (G, H) RT-qPCR analyses of liver Mup20 and Mups expression in *Cbs*^−/−^ and *Cbs*^+/−^ mice; 6-10 mice/group. Relative levels of Mup20 mRNA (G) and Mups mRNA (H) are shown. An asterisk indicates a significant difference.

Quantitative analysis showed that Mup20 protein levels were similar in the livers of 8-week-old *Cbs*^−/−^ females, *Cbs*^−/−^ males, and *Cbs*^+/−^ males **(Figure 3C)** as it was in the plasma of these mice **(Figure 3E).** In contrast, Mups proteins levels were significantly reduced in *Cbs*^−/−^ females and *Cbs*^−/−^ males, compared to corresponding *Cbs*^+/−^ animals, both in the liver **(Figure 3D)** and plasma **(Figure 3F).** Similar effects of *Cbs* genotype on Mup levels were observed in 1-year-old mice **(Figure 3C-F).**

### *Cbs*^−/−^ genotype affects transcription of the *Mup* genes in the liver

To determine whether *Cbs*^−/−^ genotype affects transcription of *Mup* genes, we quantified liver Mup mRNAs using RT-qPCR. We found that liver Mup20 mRNA, undetectable in *Cbs*^+/−^ females, was present in *Cbs*^−/−^ females **(Figure 3G**), reflecting levels of Mup20 protein **(Figure 3C)** in these mice.

In *Cbs*^−/−^ males, liver Mup20 mRNA levels were reduced (in 8-week-old animals) or unaffected (in 1-year-old animals), compared to *Cbs*^+/−^ males **(Figure 3G).** This pattern of Mup20 mRNA expression seems not reflect the levels of Mup20 protein **(Figure 3C),** suggesting a more complex regulation of Mup20 expression in males.

Other Mups mRNA levels were significantly reduced in *Cbs*^−/−^ females and *Cbs*^−/−^ males compared to corresponding *Cbs*^+/−^ mice, similarly in 8-week-old and 1-year-old animals **(Figure 3H).** This pattern of Mups mRNAs expression reflects the patern of expression of Mups protein **(Figure 3C)** in these mice. Altogether, these findings indicate that *Cbs* genotype affects transcriptional regulation of Mup expression.

### *Cbs*^−/−^ genotype downregulates the expression of Zhx2, growth hormone and sex hormone receptors in the liver

Mup expression in the liver is known to be regulated by Zhx2 transcription factor [30], growth hormone [31], and sex hormones [20]. To determine whether effects of *Cbs*^−/−^ genotype on Mup expression are mediated by Zhx2, we quantified Zhx2 protein by Western blotting **(Figure 4A)** and Zhx2 mRNA by RT-qPCR **(Figure 4B)** in livers of 8-week-old *Cbs*^−/−^ and *Cbs*^+/−^ mice. We found that Zhx2 protein was significantly reduced in *Cbs*^−/−^ mice compared to *Cbs*^+/−^ animals, with about 2-fold reduction both in males and in females **(Figure 4B).** A similar about 2-fold reduction of Zhx2 mRNA was observed in *Cbs*^−/−^ compared to *Cbs*^+/−^ mice, independently of sex **(Figure 4C).**

**Figure 4.**
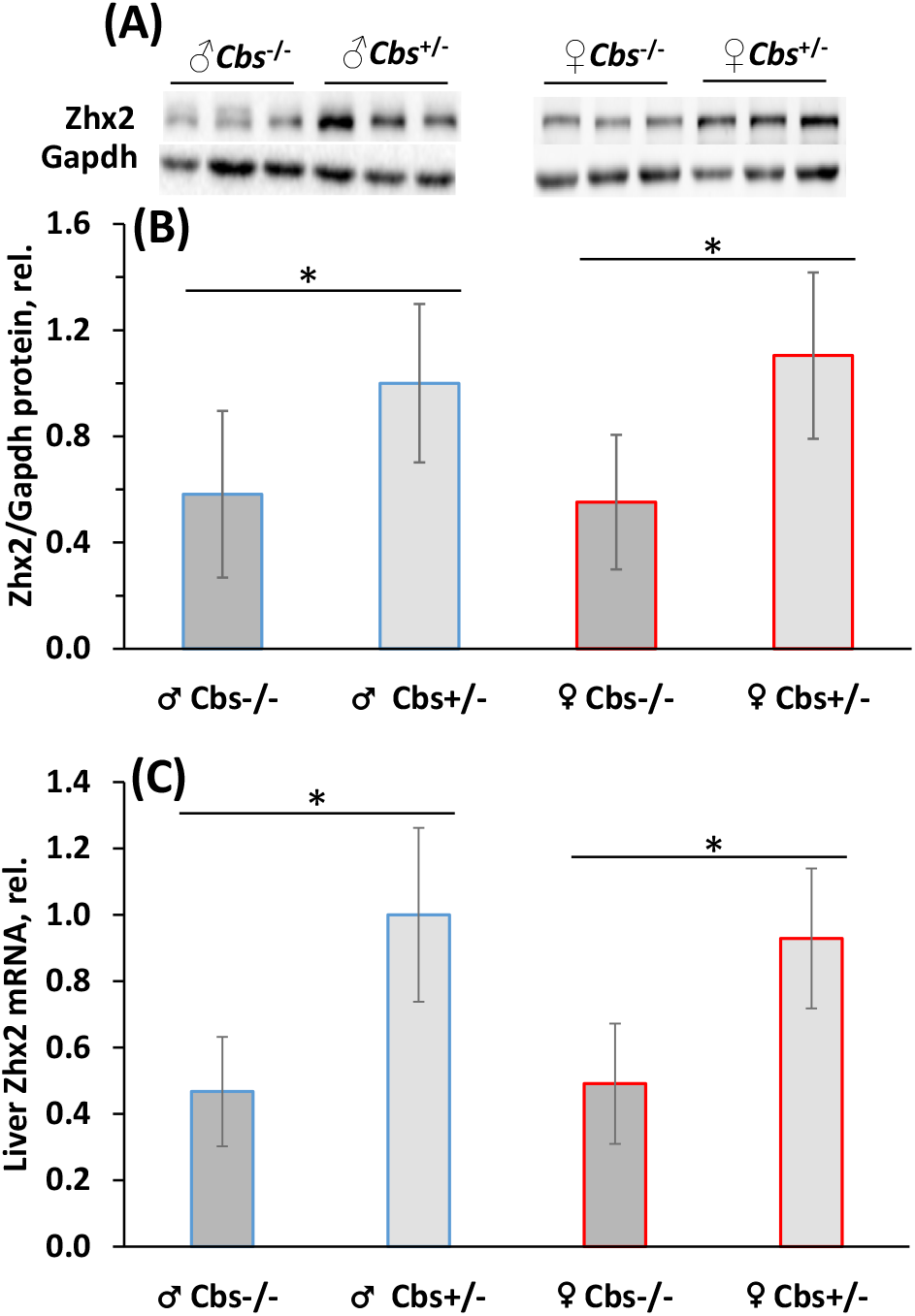
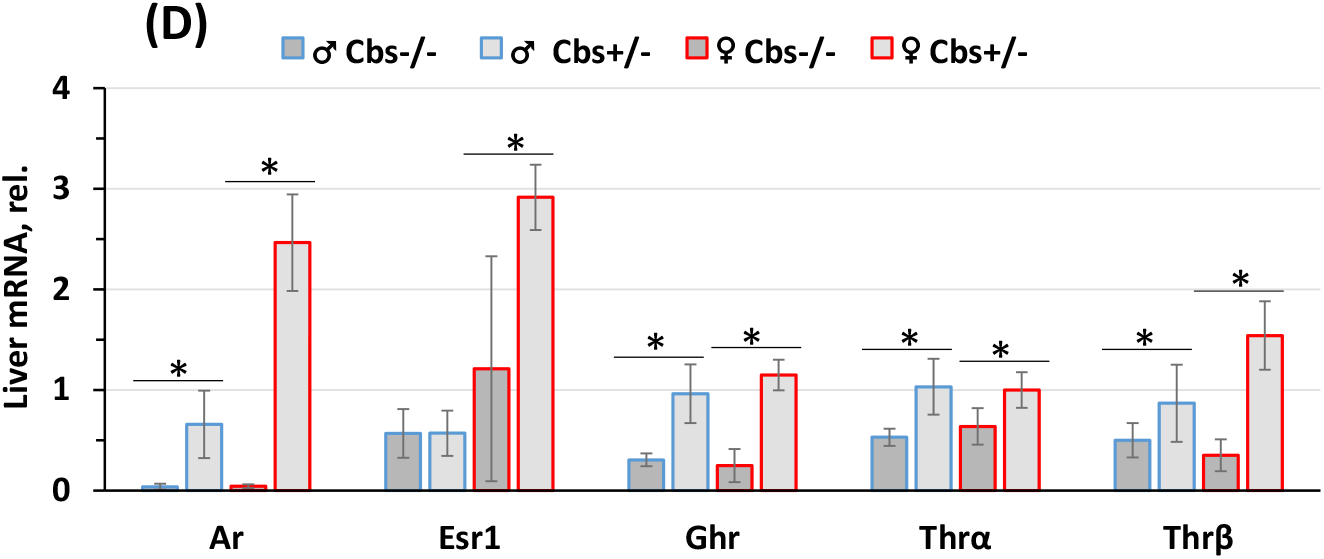
Reduced expression of Zhx2 and hormone receptors in livers of *Cbs*^−/−^ *vs. Cbs*^+/-^ mice. (A), representative Western blots of liver Zhx2. (B) Quantification of Zhx2 Western blots. (C) Quantification of liver Zhx2 mRNA by RT-qPCR. (D) Quantification of mRNAs for androgen receptor (Ar), estrogen receptor 1 (Esr1), growth hormone receptor (Ghr), thyroid hormone receptor α (Trhα), and thyroid hormone receptor β (Trhβ). Eight-week-old □ *Cbs*^−/−^, □ *Cbs^+/-^*, □ *Cbs*^−/−^, and □ *Cbs^+/-^* mice, 4-10/group, were examined. An asterisk indicates a significant difference.

To determine whether effects of *Cbs*^−/−^ genotype on Mup expression are mediated by hormones, we quantified by RT-qPCR growth hormone receptor (Ghr) mRNA and sex hormone receptors mRNAs in livers of 8-week-old *Cbs*^−/−^ and *Cbs*^+/−^ mice. We found that mRNAs for Ghr, androgen receptor (Ar), estrogen receptor (Ers), thyrotropin-releasing hormone-α (Thrα) and Thrβ were significantly reduced in *Cbs*^−/−^ mice compared to *Cbs*^+/−^ animals, with about 2-15-fold reduction in males and 2-60-fold reduction in females **(Figure 4D),** with the exception of Esr1, which was not affected by *Cbs* genotype in males. Similar results were obtained with 1-year-old mice **(Figure S2).**

### Behavioral testing

Male mice defend their territory from other males by countermarking their urine marks. Males are also attracted to female scent. We examined how *Cbs* genotype of urine donors affects these male responses. We found that male *Cbs*^−/−^ urine did not elicit a significant response in *Cbs*^+/−^ male mice. In contrast, male *Cbs*^−/−^ urine elicited a significant response from male *Cbs*^+/−^ mice, which deposited significantly more of their urine marks in the vicinity of unfamiliar male *Cbs*^+/−^ urine **(Figure 5A).** However, *Cbs*^+/−^ males were similarly attracted to female *Cbs*^−/−^ and female *Cbs*^+/−^ urines, equally occupying the test and control areas of the cage **(Figure 5B).**

**Figure 5.**
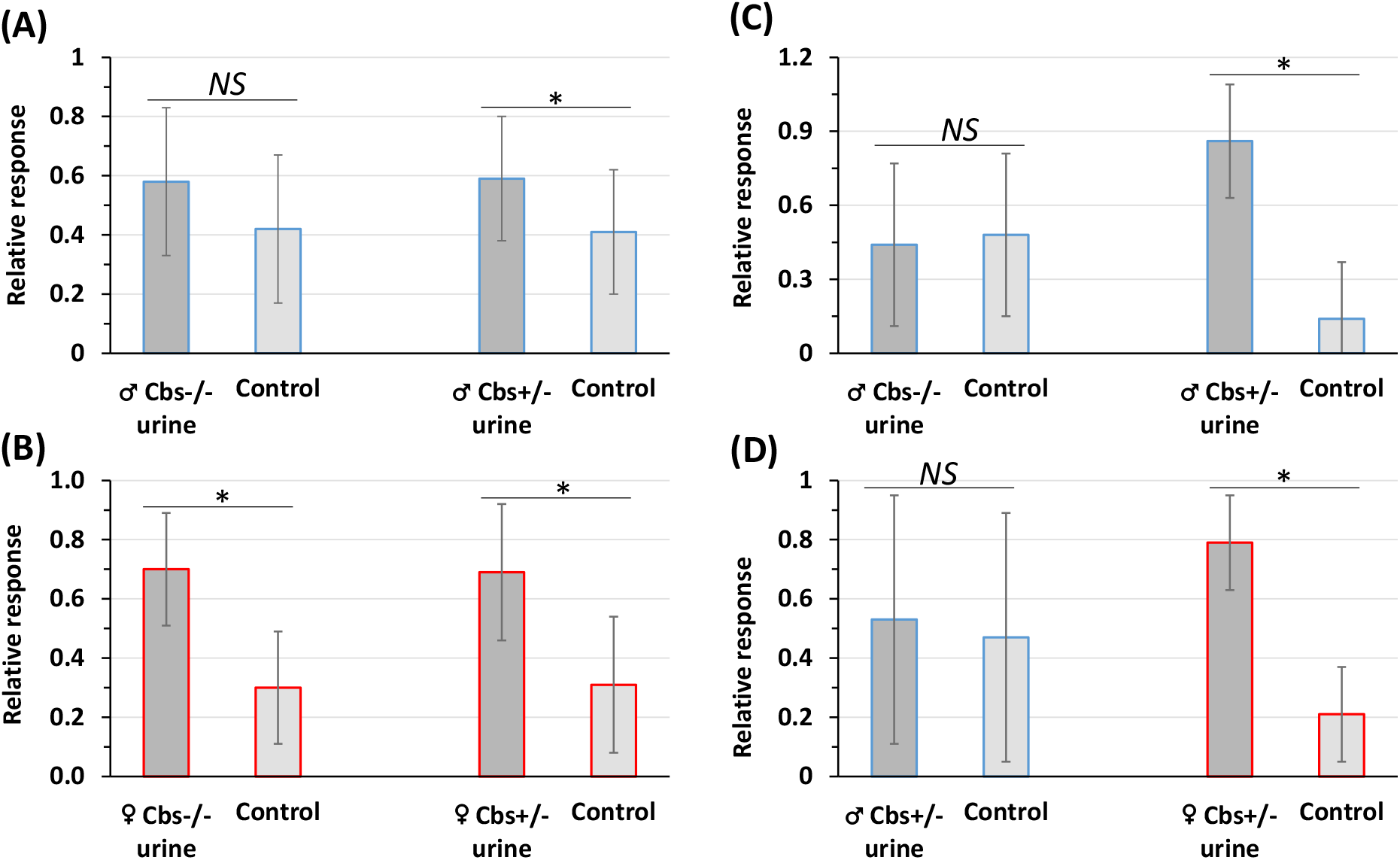
Behavioral testing of mouse responses to urines from *Cbs*^−/−^ and *Cbs*^+/−^ mice. Panels (A) and (B) illustrate responses of □ *Cbs*^+/−^ mice to urines from □ *Cbs*^−/−^ and □ *Cbs*^+/−^ mice (A), □ *Cbs*^−/−^ and □ *Cbs*^+/−^ mice (B). Panel C shows responses of □ *Cbs*^+/−^ mice to urines from □ *Cbs*^−/−^ and □ *Cbs*^+/−^ mice. Panel (D) illustrates responses of □ *Cbs*^−/−^ mice to urines from □ *Cbs*^+/−^ and □ *Cbs*^+/−^ mice. ‘Relative response’ refers to a fraction of the indicated area (urine or control) occupied by tested mice. An asterisk ‘*’ indicates a significant difference. *NS,* nor significant.

As female mice are attracted to male scent, we examined how *Cbs* genotype of urine donors affects this female response. We found that male *Cbs*^−/−^ urine was not attractive to *Cbs*^+/−^ females as shown by their equal occupation of the test and control areas of the cage **(Figure 5C).** However, a positive control experiment showed that *Cbs*^+/−^ male urine was attractive to *Cbs*^+/−^ females, which preferred to occupy the area in the vicinity of *Cbs*^+/−^ male urine **(Figure 5C).**

We also tested responses of *Cbs*^−/−^ males to control male and female urines. We found that *Cbs*^−/−^ males, did not countermark urine of control *Cbs*^+/−^ males but were still attracted to the scent of urine from control *Cbs*^+/−^ females **(Figure 5D).**

## Discussion

We studied how Mup biogenesis and sexual signaling are affected by the inactivation of the *Cbs* gene, a metabolic gene associated with female infertility, in mice. We found that sexually dimorphic Mup expression was dysregulated in *Cbs*^−/−^ mice compared to *Cbs*^+/−^ controls and that the effects of *Cbs* genotype were Mup specie- and sex-dependent. Notably, in *Cbs*^−/−^ females the expression of male-specific 18,893 Da Mup20 was derepressed while 18,709 Da specie was significantly downregulated **(Figure 2F).** In contrast, in *Cbs*^−/−^ males 18,893 Da Mup20 and 18,709 Da Mup specie were signifcantly upregulated while 18,645 Da Mup specie was significantly downregulated **(Figure 2G).** The 18,645 Da Mup specie was essentially absent in females both in *Cbs*^+/−^ and *Cbs*^−/−^ mice. The 18,693 Da Mup specie was significantly downregulated in *Cbs*^−/−^ females and males. We also found that the dysregulation of Mup biogenesis in *Cbs*^−/−^ mice reduced the quality of the *Cbs*^−/−^ urine as a source of sexual signals and impaired their sexual responses.

Our findings that *Cbs* genotype similarly affected Mup abundance in the urine **(Figure 2F, G),** plasma **(Figure 3B, E, F**), and the liver **(Figure 3A, C, D),** which is the major Mup-producing organ, indicate that *Cbs* genotype influences Mup biogenesis in the liver but does not seem to affect Mup transit from the liver to plasma to urine. Measurements of Mup protein and Mup mRNA in the liver indicate that *Cbs* genotype affected Mup biogenesis at the level of transcription in the liver **(Figure 3G, H).**

Mup levels are known to decrease with increasing mouse age [19]. However, we found that effects of age on Mup were abrogated in *Cbs*^−/−^ mice, both in males and in females. Specifically, Mup levels decreased with age in *Cbs*^+/−^ mice of both sexes as expected but the age-Mup relationship was essentially absent in *Cbs*^−/−^ mice **(Figure 1B, C).**

Growth hormone (Gh) plays an important role in the regulation of Mup transcription and sex differences in Mups expression by controling chromatin accessibility in the liver [20, 31]. Pituitary Gh secretion controls sex differences in *Mup* gene expression *via* the JAK2-STAT5 signalling pathway, which begins with Gh binding to Gh receptors (Ghr) on target cells in the liver. The Gh-Ghr binding activates the transcription factor STAT5, which enters the nucleus and initiates Mup gene transcription. Our fidings that Ghr mRNA expression was reduced 3-4-fold in *Cbs*^−/−^ males and females compared to *Cbs^+/-^* animals **(Figure 4D),** suggests that the reduction of Mups levels in mice *Cbs*^−/−^ is mediated by reduced Ghr signaling.

The transcription factor Zhx2 has been recently identified as a positive regulator that binds to and activates hepatic Mup expression [30]. In the present study we found that hepatic Zhx2 protein and mRNA expression was significantly reduced in *Cbs*^−/−^ mice and that the reduction of Zhx2 expression was similar in males and female. These findings suggest that the effects of the *Cbs*^−/−^ genotype on Zhx2 expression are transcriptional. These findings also suggest that Zhx2 regulates *Cbs* genotype-dependent Mups expression and explains the reduction of Mups levels in *Cbs*^−/−^ mice. However, our findings suggest that Zhx2 is unlikely to have any significant effect on Mup20 expression in *Cbs*^−/−^ mice.

It was not knwn whether Zhx2 affects variation in Mup levels between males and females or within the sexes. Our findings that Zhx2 expression is similarly reduced by the *Cbs*^−/−^ genotype in females and males suggest that Zhx2 is unlikely to contribute to sexual dimorphism in Mup profiles.

Several Mup genes, including Mup20 and other Mups show differential responsivness to Zhx2 [30]. In Balb/cJ male mice, deletion of Zhx2 reduced hepatic Mup20 mRNA levels to 41% of values for the hepatocyte-specific deletion (*Zhx2^Δhep^*) and 2% for whole-body deletion (*Zhx2^KO^*). Mup7, Mup10, and Mup19 mRNAs were reduced to 21% in *Zhx2^Δhep^* and 6% *Zhx2^KO^*, while Mup3 mRNA was not changed in *Zhx2^Δhep^* and reduced to 34% in *Zhx2^KO^* male mice. Western blotting with a pan-Mup antiserum showed that Mup proteins were reduced in livers of *Zbx2^Δhep^* and *Zhx2^KO^* male mice. Surprisingly, the pan-Mup antiserum showed only one band on the liver protein Western (Fig. 3 in ref [30]). In contrast, two bands showed on the urinary protein Western the pan-Mup antiserum: a major slower migrating band of the same molecular weight as the band on the liver Western and a minor faster migrating protein band corresponding to Mup20 [32] (mistakenly labeled as Mup17 in ref. [30]). Notably, only the abundant Mup protein band was responsive to changes in Zhx2 while the minor Mup20 band did not seem to respond to Zhx2 (Fig. 3 in ref [30]).

Available evidence shows that Mups influence male odor and sexual behavior as pheromones. For example, the male-specific Mup20 is attractive to females [24]. However, other Mups may influence attractiveness of the male urine scent marks to females. This possibility is supported by our findings that females were attracted to *Cbs*^+/−^ male urine but not to urine from *Cbs*^−/−^ male **(Figure 5C),** which had significantly more Mup20 and significantly reduced 18,645 Da and 18693 Da Mup species compared with *Cbs*^+/−^ male urine **(Figure 2G).**

In contrast, no studies have shown that Mups influence female odor and sexual behavior. In the present study, we found that levels of individual Mup species differ greatly between *Cbs*^−/−^ and *Cbs*^+/−^ females **(Figure 2F).** Yet, these female Mup differences seem not to bother males, which were attracted to both *Cbs*^−/−^ and *Cbs*^+/−^ female urines **(Figure 5B).** These findings suggest that female odor and sexual attractivness to males may not be influenced by female’s Mup makeup.

*We* found that althoug Mup20 is highly expressed in *Cbs*^−/−^ males, their urine does not invoke any sexual interest from control females nor males **(Figure 5D).** These findings suggest that Mup20 may not be not sufficient to assure the maleness phenotype and that other factors affected by the *Cbs*^−/−^ genotype are important as well.

In summary, our data provide evidence that Cbs is a regulator important for maintaining sexually dimorphic Mup expression in the mouse liver. An absence of the *Cbs* gene dysregulates Mup expression and impairs sexular responses in *Cbs*^−/−^ males and females. These effects are mediated by the downregulation of transcriptional factors Zhx2 and Ghr in *Cbs*^−/−^ mice. However, the mechanism of derepresion of Mup20 expression in female *Cbs*^−/−^ mice remains to be examined.

## ABBREVIATIONS

Ar: androgen receptor;
Cbs: cystathionine β-synthase;
Esrl: estrogen receptor 1;
Gapdh: glyceraldehyde-3-phosphate dehydrogenase;
Ghr: growth hormone receptor;
Hey: homocysteine;
HHcy: hyperhomocysteinemia;
Mup: major urinary protein;
RT-qPCR: reverse transcription-quantitative polymerase chain reaction;
SDS-PAGE: sodium dodecyl sulphate polyacrylamide gel electrophoresis;
Thrα: thyrotropin-releasing hormone alpha;
Tf: transferrin;
Zhx2: zink fingers and homeobox 2;

## ACKNOWLEDGMENTS

Supported in part by grants 2014/15/N/NZ5/01646, 2016/23/B/NZ5/00573, 2018/29/B/NZ4/00771, and 2019/33/B/NZ4/01760 from the National Science Center, Poland.

## AUTHOR CONTRIBUTION

E. Bretes performed and analyzed the biochemical experiments and contributed to writing the first draft of the paper; J. Wróblewski performed the biochemical experimnets and contributed to breeding the mice; M. Wyszczelska-Rokiel performed the behavioral experiments and analyzed the data; H. Jakubowski conceived the idea for the project, designed the study, bred the mice, collected mouse urine, plasma and tissue samples, performed the behavioral experiments, analyzed data, and wrote the paper.

## Data Availability Statement

The data that support the findings of this study are available in the methods and/or supplementary material of this article.

## Supplementary material

**Figure S1.**
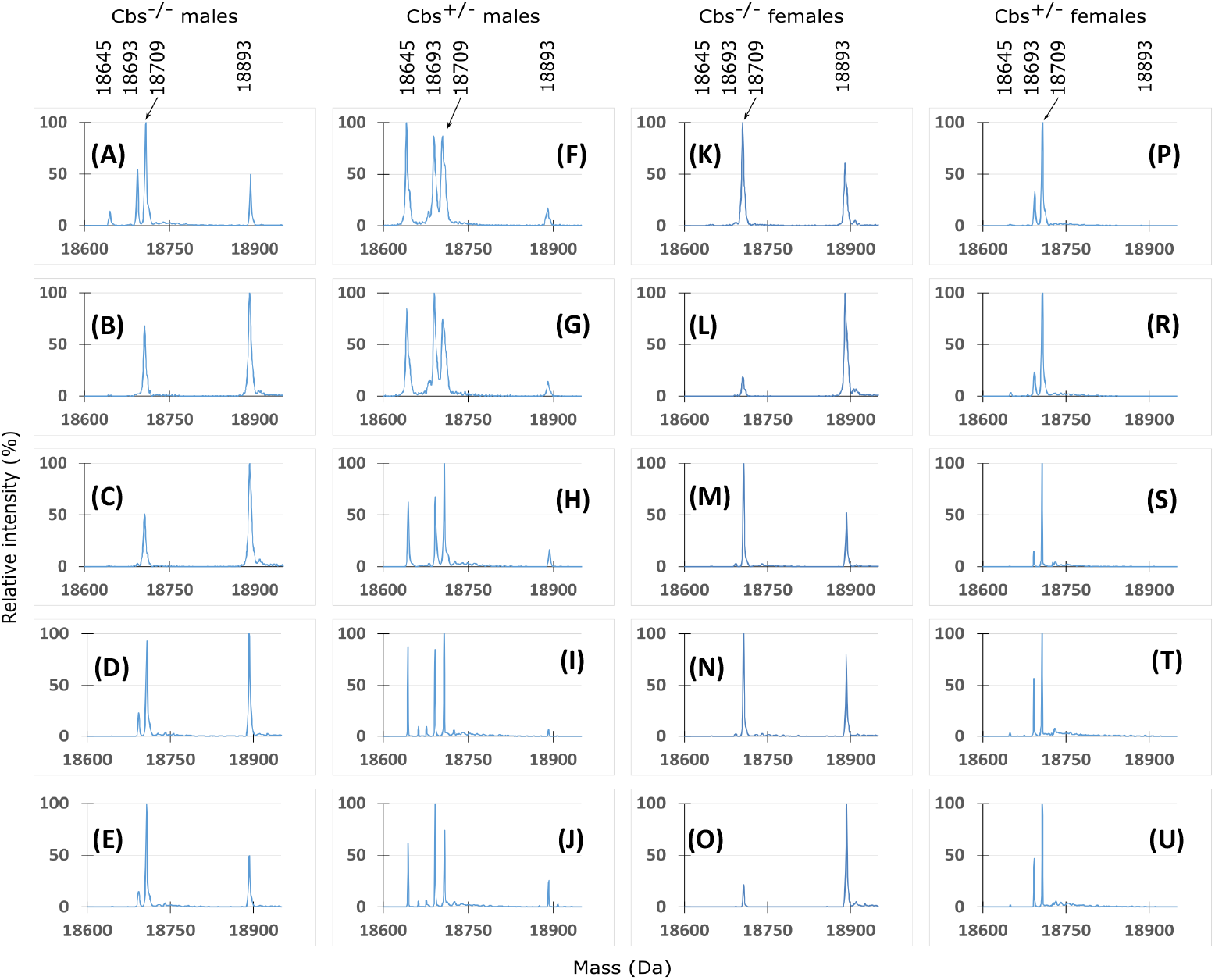
ESI-MS profiles of individual Mup species in *Cbs*^−/−^ and *Cbs*^+/−^ mice. Urines from □ *Cbs*^−/−^ mice (A-E), □ *Cbs*^+/−^ mice (F-J), □ *Cbs*^−/−^ mice (K-O), and □ *Cbs*^+/−^ mice (P-U) were analyzed (5 animals/group). Molecular weights (Da) are indicated above each set of peaks. Male Mup20 of 18,893 Da, absent is control □ *Cbs*^+/−^ mice, is found in urine from each □ *Cbs*^−/−^ mice.

**Figure S2.**
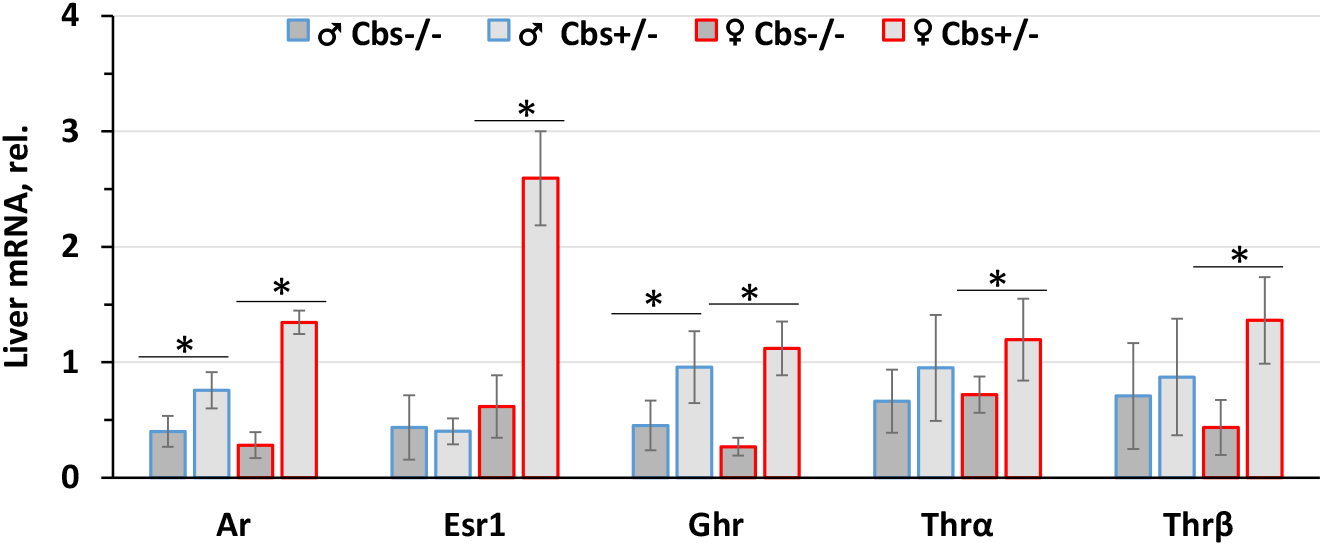
Reduced expression of sex hormone receptors in livers of old *Cbs*^−/−^ vs. *Cbs*^+/−^ mice. mRNAs for androgen receptor (AR), estrogen receptor 1 (Esr1), growth hormone receptor (GRH), thyroid hormone receptor α (TRHα), and thyroid hormone receptor β (TRHβ) were quantified in livers of 1-year-old *Cbs*^−/−^ and control *Cbs*^+/−^ mice of both sexes (n = 6/group). An asterisk indicates a significant difference.

**Table S1.**
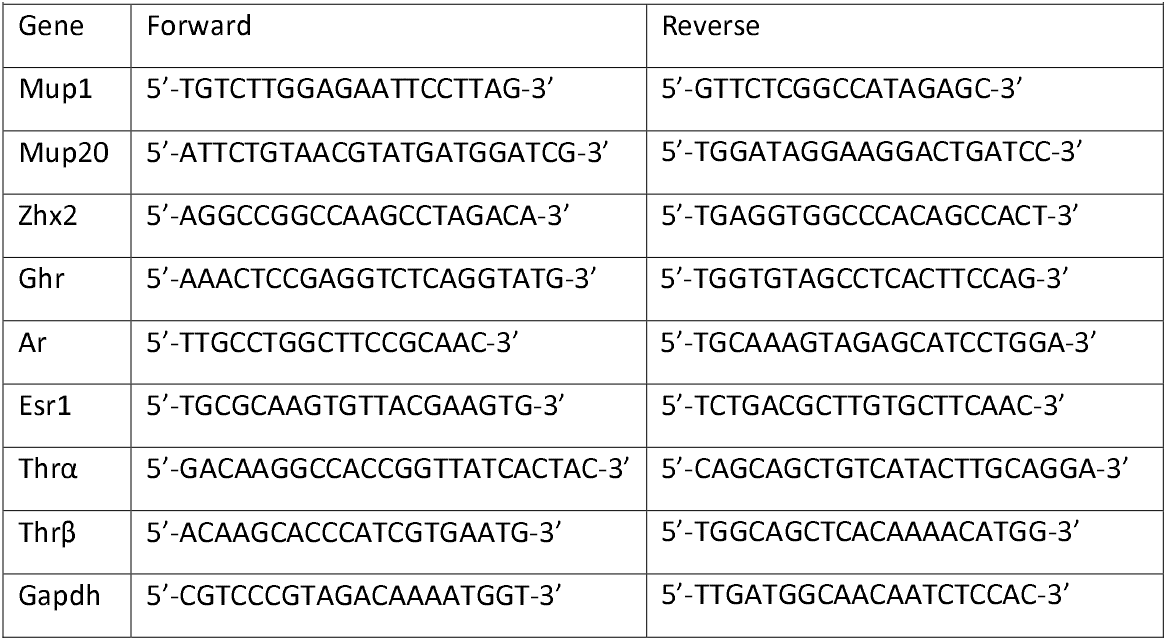
Oligonucleotide sequences used in the present study.

