## Supplementary material for "Cystathionine β-synthase gene inactivation dysregulates major urinary protein biogenesis and impairs sexual signaling in mice": Suppl Info

Cystationine β-synthase gene inactivation derepresses female Mup20 expression and impairs sexual signaling in mice

**Supplemental Information**

**Figure S1.** ESI-MS profiles of individual Mup species in *Cbs*^-/-^ and *Cbs*^+/-^ mice. Urines from ♂ *Cbs*^-/-^ mice (A-E), ♂ *Cbs^+/-^* mice (F-J), ♀ *Cbs^-/-^* mice (K-O), and ♀ *Cbs^+/-^* mice (P-U) were analyzed (5 animals/group). Molecular weights (Da) are indicated above each set of peaks. Male Mup20 of 18,893 Da, absent is control ♀ *Cbs^+/-^* mice, is found in urine from each ♀ *Cbs^-/-^* mice.

**Figure S2.** Reduced expression of sex hormone receptors in livers of old *Cbs^-/-^* vs. *Cbs^+/-^* mice. mRNAs for androgen receptor (AR), estrogen receptor 1 (Esr1), growth hormone receptor (GRH), thyroid hormone receptor α (TRHα), and thyroid hormone receptor β (TRHβ) were quantified in livers of 1-year-old *Cbs*^-/-^ and control *Cbs^+/-^* mice of both sexes (n = 6/group).

**Table S1.** Oligonucleotide sequences used in the present study

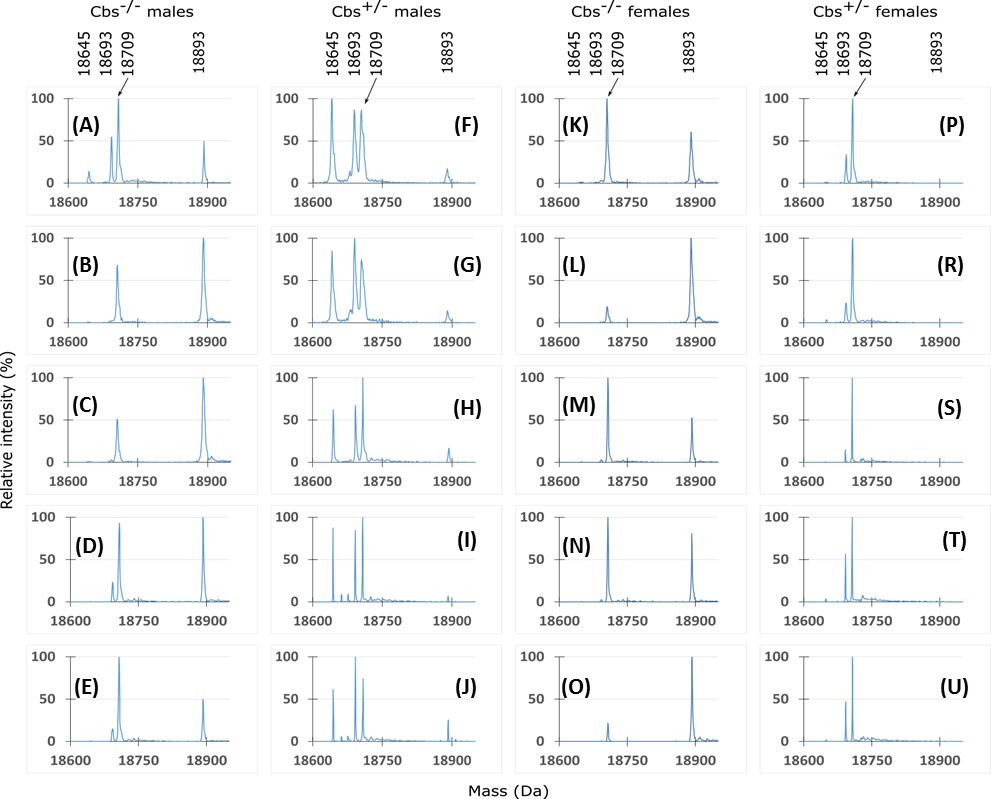

**Figure S1**

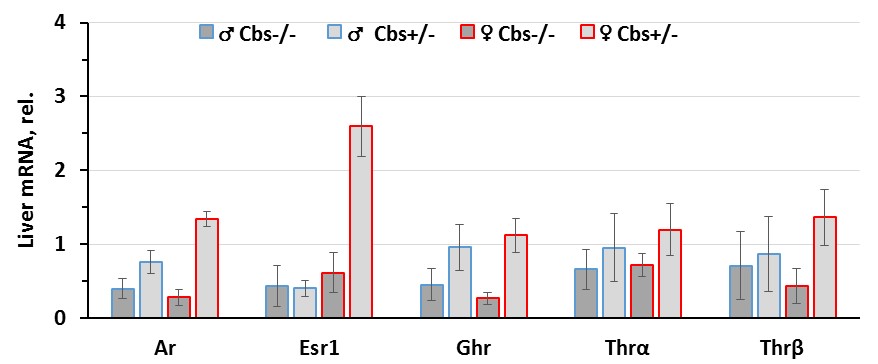

**Figure S2**

| **Table S1.** Oligonucleotide sequences used in the present study | | |
| --- | --- | --- |
| Gene | Forward | Reverse |
| Mup1 | 5’-TGTCTTGGAGAATTCCTTAG-3’ | 5’-GTTCTCGGCCATAGAGC-3’ |
| Mup20 | 5’-ATTCTGTAACGTATGATGGATCG-3’ | 5’-TGGATAGGAAGGACTGATCC-3’ |
| Zhx2 | 5’-AGGCCGGCCAAGCCTAGACA-3’ | 5’-TGAGGTGGCCCACAGCCACT-3’ |
| Ghr | 5’-AAACTCCGAGGTCTCAGGTATG-3’ | 5’-TGGTGTAGCCTCACTTCCAG-3’ |
| Ar | 5’-TTGCCTGGCTTCCGCAAC-3’ | 5’-TGCAAAGTAGAGCATCCTGGA-3’ |
| Esr1 | 5’-TGCGCAAGTGTTACGAAGTG-3’ | 5’-TCTGACGCTTGTGCTTCAAC-3’ |
| Thrα | 5’-GACAAGGCCACCGGTTATCACTAC-3’ | 5’-CAGCAGCTGTCATACTTGCAGGA-3’ |
| Thrβ | 5’-ACAAGCACCCATCGTGAATG-3’ | 5’-TGGCAGCTCACAAAACATGG-3’ |
| Gapdh | 5’-CGTCCCGTAGACAAAATGGT-3’ | 5’-TTGATGGCAACAATCTCCAC-3’ |
